# Evaluating the efficiency and accuracy of a commercial test for estimating genetic risk of bovine congestive heart failure

**DOI:** 10.1101/2023.04.24.536588

**Authors:** Jaden M. Carlson, Michael P. Heaton, Nathan Allison, Alyssa Hangman, Dustin Petrik, Heather Piscatelli, Brian L. Vander Ley

**Author notes:** **Correspondence:** Brian L. Vander Ley.

## Abstract

**Background:** Bovine congestive heart failure (BCHF) is a significant cause of death in feedlot cattle in the Western Great Plains of North America. Single nucleotide polymorphisms (SNPs) in the *ARRDC3* and *NFIA* genes have been previously associated with BCHF and genetic tests can classify animals by their risk for disease. Here, our aims were to evaluate the efficiency (genotypes obtained / samples tested) of a rapid DNA extraction kit and the accuracy of a 2-SNP assay for BCHF risk.

**Methods:** Skin biopsies from 100 cattle were randomized and extracted with a proprietary rapid DNA extraction kit. A custom duplex, combined sequence amplification and nucleotide detection (C-SAND) assay was developed and run once on a commercial thermocycling machine to determine the genotypes. Both the rapidly extracted DNA and highly purified reference DNA from the same individuals were genotyped with the 2-SNP assay by operators blinded to the sample identity. The C-SAND genotypes were compared to known genotypes derived from a bead array assay. *A priori* standards for missing and incorrect genotypes were set at less than 3% and 1%, respectively.

**Results:** When using reference DNA samples, there were no missing and no incorrect C-SAND-derived genotypes, meeting the *a priori* standards. When DNA samples from the rapid extraction kit were used, genotypes were not determined for 5% of the samples. Of the 95 samples successfully extracted, there were 0% and 3% incorrect genotypes for the respective *ARRDC3* and *NFIA* SNPs.

**Conclusions:** This duplex C-SAND assay and thermocycling machine combination were efficient and accurate when reference DNA was used, meeting *a priori* standards. Although the reduced efficiency of the rapid extraction kit can be overcome by repeated testing, increased genotype errors present an important issue. Despite these challenges, this rapid extraction kit and assay can be a reasonable tool for producers to select animals with reduced BCHF risk.

## Introduction

Bovine congestive heart failure (BCHF) is an emerging cause of production and death loss for cattle late in the feeding period at low to moderate elevations (Jensen et al., 1976; Johnson et al., 2021; 2022; Neary et al., 2016). Although details of the pathogenesis are complex and not well understood, BCHF is generally attributed to ‘*cor pulmonale*’ which is the enlargement of the right heart as a result of disease of the lungs or pulmonary blood vessels. This eventually leads to right ventricular overload and ultimately heart failure (Figure 1). Affected cattle have various symptoms, including biventricular fibrosis, cardiac adipose depots, coronary artery injury, and pulmonary venous remodeling (Krafsur et al., 2019;Moxley et al., 2019). Some evidence suggests that hypobaric hypoxia and left heart dysfunction may both play a role in disease development (Krafsur et al., 2019). However, the disease pathogenesis of BCHF in feedlot cattle maintained at low to moderate elevations remains unclear. Regardless, losses due to BCHF in severely affected operations can rival those caused by bovine respiratory disease (Heaton et al., 2019). Thus, reducing the impact of BCHF is a high priority for the beef cattle industry.

**Figure 1.**
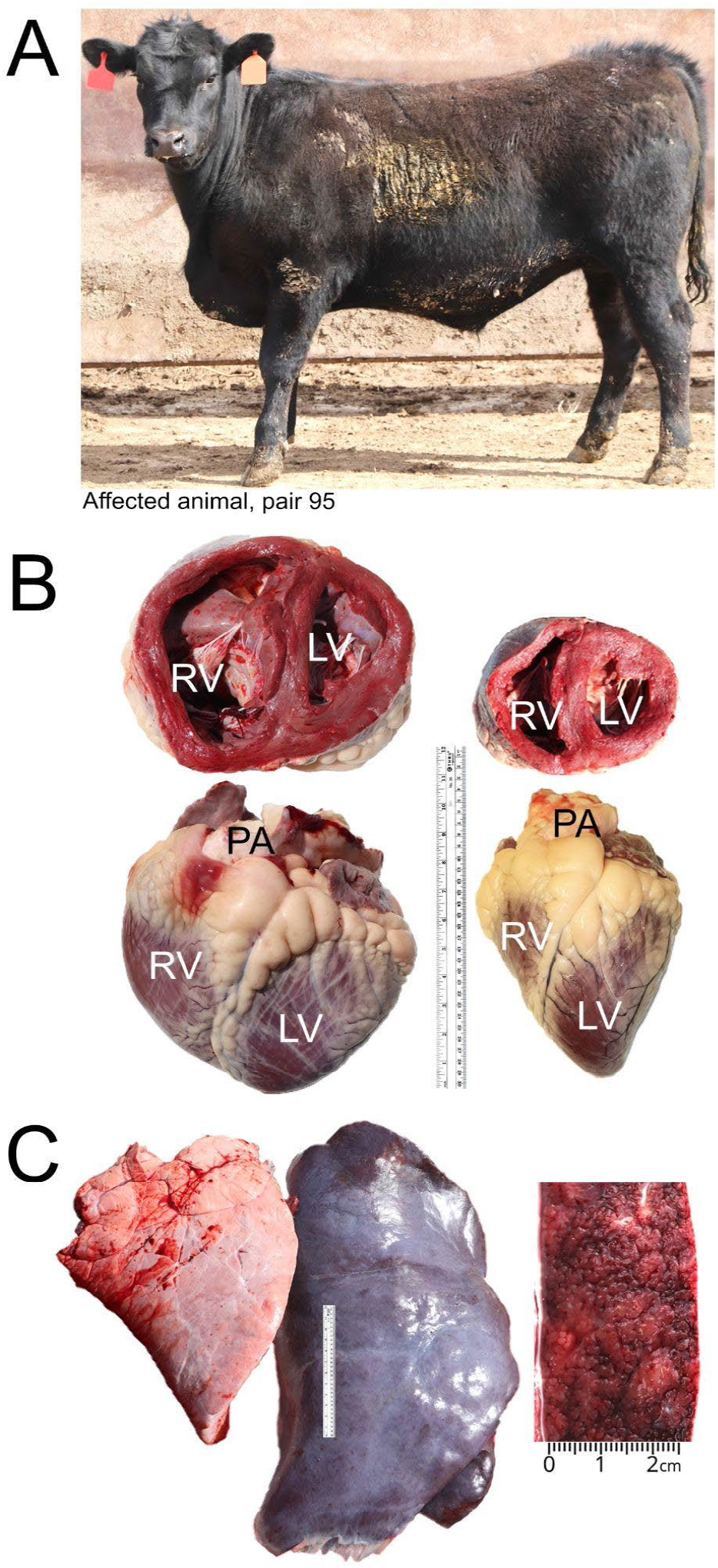
Clinical presentation of BCHF and gross morphological features of heart and liver in a feedlot steer. Panel A, the clinically affected animal from case-control pair number 95 fattened at 1,200 m displaying ventral edema and a drop in the withers due to fluid accumulation in the thorax which causes the shoulders to be pushed outward (Heaton et al., 2019). Panel B, gross heart morphology of affected case (left) compared with a normal heart of a fattened Angus heifer at 578 m (right). Abbreviations: RV, right ventricle; LV, left ventricle; PA, pulmonary artery. Panel C, lungs (left), enlarged encapsulated liver (center), with ‘nutmeg liver’ appearance in cross section due to hepatic venous congestion (right).

Although the etiology of BCHF is unknown, several environmental and management risk factors have been proposed based on associations in retrospective analyses and similarities with heart failure in chickens and humans. These include prior exposure to hypoxic conditions at high elevation, rapid growth rate, intensive or prolonged fattening period, and sex (Jensen et al., 1976; Johnson et al., 2021; 2022; Krafsur et al., 2019; Neary et al., 2016). Despite these and other epidemiological studies, there is a lack of consensus on the non-genetic risk factors associated with BCHF in cattle at low to moderate elevations. However, multiple genetic risk factors associated with heart failure in cattle have been reported. A candidate gene (*EPAS1*) variant was reported to be associated with pulmonary hypertension in Angus cattle at high elevations (Newman et al., 2015). This was a double missense variant haplotype (T606/S610) in the gene encoding the hypoxia inducible factor two alpha (HIF2a) protein. Four additional haplotype variants of *EPAS1* were also reported from whole genome sequence (WGS) analysis of a diverse panel of 96 bulls representing 19 common beef breeds (Heaton et al., 2016). However, there was no association between any *EPAS1* haplotypes and BCHF at moderate elevations with 204 matched case-control pairs where the case had clinical, end stage BCHF (Heaton et al., 2019). Subsequently, a genome wide association study (GWAS) with the same 204 matched pairs and 560,000 single nucleotide polymorphism (SNP) markers identified a significant association between BCHF and linked variants in the arrestin domain-containing protein 3 (*ARRDC3*) gene and nuclear factor IA (*NFIA*) gene (Heaton et al., 2022). Compared to their pen mates without homozygous risk alleles, cattle with homozygous risk alleles at either gene were 8-fold more likely to have BCHF. Additionally, cattle with homozygous risk alleles at both genes were 28-fold more likely to have BCHF. Successful use of these BCHF-associated DNA markers will require accurate genotyping tests to assist cattle producers in managing severely affected herds.

The availability of commercial genetic tests for *ARRDC3* and *NFIA* variants associated with BCHF is currently limited. An assortment of custom proprietary bead chip arrays are available; however, the critical SNP marker data is not accessible to most producers, and some bead arrays may not contain the BCHF SNP markers. Further complicating the use of custom bead chip arrays is the issue of data ownership when a proprietary test is provided as part of a breed association requirement. Another technology currently available is whole genome sequencing (WGS) (e.g., 15x coverage) which is a custom service that is cost-prohibitive, slow, and requires data-intensive tools and expertise out of reach for most cattle producers. Low pass genome sequencing (e.g., 0.5x coverage) combined with imputation represents a potential solution; however, establishing the best practices pipeline to achieve the desired accuracy is not trivial (Buckley et al., 2022). Regardless of platform, and assuming a genotype scoring accuracy of 99%, producers may still wish to have two independent tests performed to avoid the unnecessary culling of valuable cattle. Thus, there is still a need for separate, custom, accurate genotyping tests to classify cattle for BCHF risk.

As part of a previously published GWAS study, a 2-SNP assay was developed to classify genetic risk for BCHF (Heaton et al., 2022). One marker was a missense variant (C182Y) in the *ARRDC3* gene (identifiers rs109901274, BovineHD0700027239, BCHF5) while the other marker was in intron 4 of the *NFIA* gene (identifiers rs133192205, BovineHD0300024308, BCHF32). Here, our aim was to assess the efficiency of a proprietary rapid DNA extraction kit and the accuracy of the genotyping platform in classifying cattle for BCHF risk with a 2-SNP test in bovine *ARRDC3* and *NFIA* genes. The metrics used were call rate (genotypes obtained out of the samples tested) and accuracy (correct genotypes out of the total genotypes). Although the *a priori* standards of 97% call rate and 99% accuracy were narrowly missed in some instances, the results indicated that this rapid extraction kit and assay can be a useful tool for identifying cattle with reduced BCHF risk so cattle producers can assess their options for classifying BCHF risk.

## Methods

### Ethics statement

The experimental design and procedures used during this research project were reviewed and approved by the Institutional Animal Care and Use Committee of the University of Nebraska-Lincoln as previously described (experimental outline numbers 139 and 1172 [Workman et al., 2016; Heaton et al., 2019]). The animals were privately owned by commercial feeding operations, and the owners and management approved the use of animals for this study. In every instance, all efforts were undertaken to reduce animal suffering. Additionally, animal welfare for these facilities was in accordance with the National Cattlemen’s Beef Association’s Beef Quality Assurance Feedyard Welfare Assessment program (http://www.bqa.org/).

### Sample selection and DNA purification

One hundred blinded, randomized reference DNAs were used from a set of 204 skin biopsy tissues that were collected and purified as previously described (Heaton et al., 2019; 2022). Briefly, small ear notches were collected, dried in granular NaCl, and stored at −20°C until use. DNA was isolated using standard procedures including RNase/protease digestion, phenol-chloroform extraction, and ethanol precipitation (Heaton et al., 2008). Purified DNAs were dissolved in a solution of 10 mM TrisCl, 1 mM EDTA (TE, pH 8.0), and stored at 4°C. Reference sample genotypes were determined previously by two independent methods: the BovineHD BeadChip array scored at GeneSeek Inc. (Lincoln, NE, USA), according to manufacturer’s instructions (Illumina, Inc., San Diego, CA, USA), and by custom assays as previously described (Heaton et al., 2019, 2022).

### Commercial genotyping assay evaluation

Tissues from the same 100 blinded, randomized reference DNAs described above were randomized again, renumbered, and used to measure the call rate and accuracy of the assays for rs109901274 and rs133192205 with the commercial platform (Figure 2). These SNPs were assigned assay identifiers BCHF5 (*ARRDC3*) and BCHF32 (*NFIA*), respectively (Table 1). The manufacturer’s assay was a combined sequence amplification and nucleotide detection (C-SAND) method based on ‘padlock’ oligonucleotide probes (MatMaCorp, Inc., Lincoln, NE, USA). Briefly, padlock probes were long (70 to 100-mer) oligonucleotides whose ends were complementary to adjacent sequences on the target genome. The padlock probes were combined with fluorescently labeled probes and isothermal rolling circle amplification technology, for the qualitative detection of alleles (Banér et al., 1998; Mohsen and Kool, 2016).

**Table 1.** Single nucleotide polymorphism (SNP) identifiers were used to design a 2-SNP assay that was evaluated for efficiency and accuracy using samples from feedlot cattle.

| Gene | Chr.<sup>a</sup> | ARS-UCD1.2 position<br>(base pair) | BeadChip array<br>identifier | rs number | Alias |
| --- | --- | --- | --- | --- | --- |
| *ARRDC3* | 7 | 90845941 | BovineHD0700027239 | rs109901274 | BCHF5 |
| *NFIA* | 3 | 84580655 | BovineHD0300024308 | rs133192205 | BCHF32 |

<sup>a</sup> Chromosome

**Figure 2.**
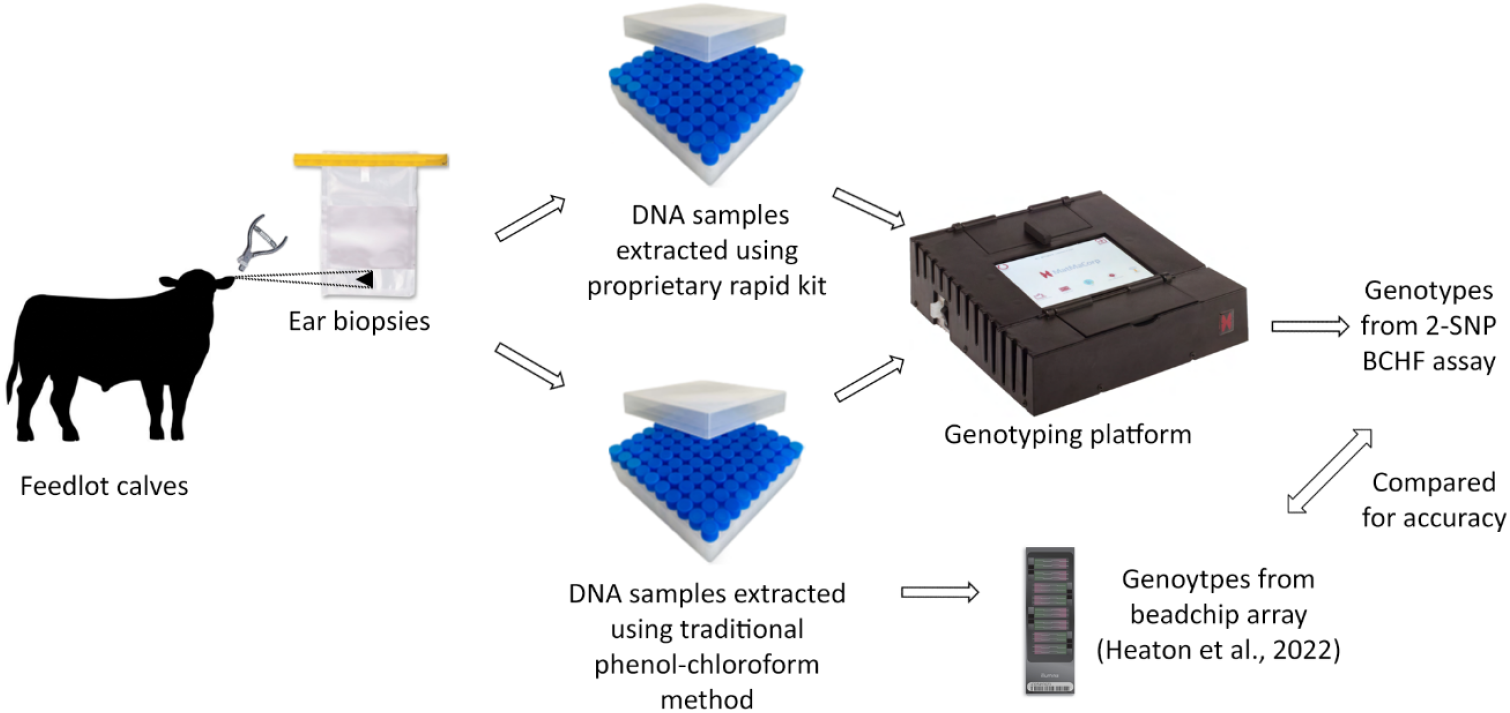
Experimental design for evaluating the call rate and accuracy of a commercial test for estimating genetic risk of BCHF. Ear biopsies from BCHF cases and their matched penmate controls were collected, isolated for reference-quality DNA, and genotyped using a bead chip array as previously described (Heaton et al., 2019; 2022). One hundred feedlot calves were randomly selected, and both the ear biopsy and reference-quality DNA was used to evaluate a proprietary rapid DNA extraction kit and commercial genotyping test for call rate and accuracy compared to the bead chip array genotypes.

The DNA from these 100 tissues was isolated with a proprietary rapid DNA extraction kit and tested by technicians at the company per the manufacturer’s instructions (MatMaCorp, Inc., Lincoln, NE, USA). A clean razor blade was used to cut a small piece of tissue from the ear notch (approximately 30 mg). An initial wash step was performed on each piece of tissue by dipping it three times in 70% ethanol. Excess ethanol was removed by blotting on a clean tissue. DNA was then isolated by following the isolation kit instructions. Once extracted, each DNA sample was genotyped with the C-SAND assay and tested on the Solas8 device (MatMaCorp, Inc., Lincoln, NE, USA) according to the manufacturer’s instructions. Following the completion of the assay, the Solas8 device automatically determined a genotype for each sample. Machine scored genotypes were recorded and compared to those made by manually analyzing the real time graphical data generated during each run. Samples with missing genotypes were not included in the accuracy calculations. Genotype results were decoded and compared to reference genotypes.

The genotype call rate and accuracy were defined in equations 1 and 2, respectively:

1. Call rate = (the number of samples with a scored genotype) / (the number of samples tested once)
2. Accuracy = (the number of correctly genotyped samples) / (the number of samples scored)

The genotype call rates and accuracies measured here were compared to standards previously established for challenging parentage exclusion testing scenarios, where only one parent is available for a given offspring (Heaton et al., 2014; 2016). A high call rate (>97%) was used to ensure that any two animals will have 94% of the same marker genotypes available for comparison for parentage exclusion. The call rate also determines the level of repeated testing required to get a result. High genotype accuracy (>99%) is critical since a single marker can exclude an animal from parentage. While the *ARRDC3* and *NFI*A markers described here were not intended for use in parentage exclusion, the conservative standards with only two non-redundant markers provides a rigorous and defined confidence level for producers.

## Results

The commercial assay had call rates and accuracies of 100% for both SNPs tested when reference DNAs were used (Table 2). When samples extracted with a proprietary rapid kit were used, the call rate dropped to 95% since five samples did not provide genotype results on the first attempt. Genotype accuracies of the remaining 95 samples were 100% and 97%, respectively, for the *ARRDC3* and *NFIA* SNPs. A second extraction was performed on the five samples with missing genotypes and all five gave correct genotypes.

**Table 2.** Call rate and accuracy of two DNA extraction methods on two single nucleotide polymorphisms associated with BCHF.

| DNA extraction type | *ARRDC3* (BCHF5) |  | *NFIA* (BCHF32) |  |
| --- | --- | --- | --- | --- |
|  | Call rate (%) | Accuracy (%) | Call rate (%) | Accuracy (%) |
| Traditional phenol-chloroform method | 100 | 100 | 100 | 100 |
| Proprietary rapid kit | 95 | 100 | 95 | 97 |

## Discussion

Here we performed a blinded evaluation of call rate and accuracy for a commercially available 2-SNP assay that determines *ARRDC3* and *NFIA* variant genotypes associated with BCHF in feedlot cattle. The assay and testing system met rigorous standards when highly purified DNA was used, and nearly met the same standards when ear biopsies were used with a proprietary rapid DNA extraction kit. A strength of this study was using hundreds of blinded samples with known genotypes from reference DNAs using the BovineHD BeadChip array. A limitation was the lack of repeated testing with new random sets of samples with known genotypes. Thus, confidence intervals were not calculated. Nevertheless, these results provide a benchmark from which to begin testing and estimate 95% success on first pass testing with this system. While the proprietary DNA extraction kit was imperfect, it’s rapidness and straightforwardness is an advantage over the lengthy and complicated process of the traditional phenol-chloroform method, especially in non-laboratory settings such as a feedlot or veterinary clinic.

This duplex C-SAND assay and thermocycling machine combination were efficient and accurate for typing *ARRDC3* and *NFIA* variants associated with BCHF when reference DNA was used, meeting *a priori* standards. When ear biopsies were extracted using the proprietary rapid DNA extraction kit, the observed reduced call rate (95%) could be overcome by retesting the 5% uncalled samples. However, the observed 3% genotype inaccuracies present an important challenge for cattle producers since tools for identifying these errors are not readily available. Despite these challenges, this proprietary rapid DNA extraction kit and assay is expected to be a useful tool for cattle producers in managing severely affected herds.

## Acknowledgements

Mention of trade names or commercial products in this publication is solely for the purpose of providing specific information and does not imply recommendation or endorsement by the U.S. Department of Agriculture. The U.S. Department of Agriculture (USDA) prohibits discrimination in all its programs and activities on the basis of race, color, national origin, age, disability, and where applicable, sex, marital status, familial status, parental status, religion, sexual orientation, genetic information, political beliefs, reprisal, or because all or part of an individual’s income is derived from any public assistance program. (Not all prohibited bases apply to all programs.) Persons with disabilities who require alternative means for communication of program information (Braille, large print, audiotape, etc.) should contact USDA’s TARGET Center at (202) 720-2600 (voice and TDD). To file a complaint of discrimination, write to USDA, Director, Office of Civil Rights, 1400 Independence Avenue, S.W., Washington, D.C. 20250-9410, or call (800) 795-3272 (voice) or (202) 720-6382 (TDD).

USDA is an equal opportunity provider and employer.

